# Engineered Oxalate decarboxylase boosts activity and stability for biological applications

**DOI:** 10.1101/2024.05.06.592502

**Authors:** Mirco Dindo, Carolina Conter, Gen-ichiro Uechi, Gioena Pampalone, Luana Ruta, Angel L. Pey, Luigia Rossi, Paola Laurino, Mauro Magnani, Barbara Cellini

## Abstract

Oxalate decarboxylase (OxDC) from *Bacillus subtilis* is a Mn-dependent hexameric enzyme which converts oxalate to carbon dioxide and formate. Recently, OxDC has attracted the interest of the scientific community, due to its biotechnological and medical applications for the treatment of hyperoxalurias, a group of pathologic conditions associated with excessive oxalate urinary excretion due to either increased endogenous production or increased exogenous absorption. The fact that OxDC displays optimum pH in the acidic range, represents a big limitation for most biotechnological applications involving processes occurring at neutral pH, where the activity and stability of the enzyme are remarkably reduced. Here, through bioinformatics-guided protein engineering, followed by combinatorial mutagenesis and analyses of activity and thermodynamic stability, we identified a double mutant of OxDC endowed with enhanced catalytic efficiency and stability under physiological conditions. The obtained engineered form of OxDC offers a potential tool for improved intestinal oxalate degradation in hyperoxaluria patients.

## Introduction

Hyperoxaluria is a pathologic condition characterized by increased urinary oxalate excretion(1–3). In humans, oxalate is a metabolic end-product that can arise from endogenous (glycolate and hydroxyproline metabolism) or exogenous (diet) sources (3–5). Interestingly, oxalate homeostasis is also affected by the gut microbiota, which can degrade oxalate and is able to modulate intestinal absorption and excretion (6). Hyperoxaluria is the main risk factor for the formation of calcium oxalate stones in the urinary tract, which can progress to nephrocalcinosis and kidney failure (1, 7). This condition can result from (i) an increased endogenous oxalate production due to genetic alterations in genes involved in glyoxylate/oxalate liver metabolism, and in this case is called Primary Hyperoxaluria (PH)(2) or (ii) alterations of the gastrointestinal tract leading to increased exogenous oxalate absorption, a condition named Secondary hyperoxaluria (SH) (1). A specific form of SH, named enteric hyperoxaluria is often observed in patient with digestive diseases and can be the result of an increased bioavailability of dietary oxalate possibly associated with an increased gastrointestinal oxalate permeability (3, 7, 8). One of the therapeutic approaches proposed for the treatment of hyperoxaluria is the oral administration of Oxalate decarboxylase (OxDC), a non-human enzyme that can degrade intestinal oxalate (9, 10). Indeed, OxDC can reduce intestinal oxalate absorption, thus counteracting the basic cause of SH(10). In addition, a low oxalate concentration in the gut can promote the intestinal excretion of plasmatic oxalate, thus possibly reducing the burden in PH patients (8). This explains why OxDC has attracted great interest from the scientific community. The initial use of OxDC focused on industrial applications(11), (i.e., preventing the formation of oxalate salt deposits in industrial processes such as papermaking and beer production). Also, the potential biomedical application of OxDC has been extended at the diagnostic level for the determination of oxalic acid concentration in food and complex biological samples such as blood and urine (12–15).

OxDC from *Bacillus subtilis* is a homohexamer (a dimer of trimers) (Figure 1) of 264 kDa which belongs to the cupin superfamily and requires Mn^2+^ and O_2_ to catalyze the conversion of oxalate to formate and CO_2_ (16). Structurally, each monomer consists of two cupin domains showing a characteristic β-sandwich structure (Figure 1A) and coordinates two Mn^2+^ ions located in the center of each cupin domain, where the metal interacts with highly conserved amino acids (16–18). OxDC cleaves the carbon-carbon bond of oxalate to CO_2_ and formate through a radical-based catalytic cycle involving electron transfer from the coordinated Mn(II) ion to the bound oxygen in domain I (N-terminal) of each monomer (16, 19). However, the identity of the catalytic active site is still highly controversial. It has been suggested that domain I is the only responsible for the enzymatic activity, whereas domain II plays mainly a structural role (16, 20). Recent studies have identified the presence of a channel for oxalate diffusion that may exist in the N-terminal domain of the monomers in an “open” or “closed” form (18). The major player involved in the conformational rearrangement of the channel is a pentapeptide loop formed by residues 161-165, generating a lid important for reaction specificity. In detail, the lid in cupin domain I contains the proton donor residue (Glu162) and isolates the active site from the solvent during catalysis(17). The oxidation state of the bound metal ion also plays a crucial role in catalyzing the OxDC reaction (21).

**Figure 1.**
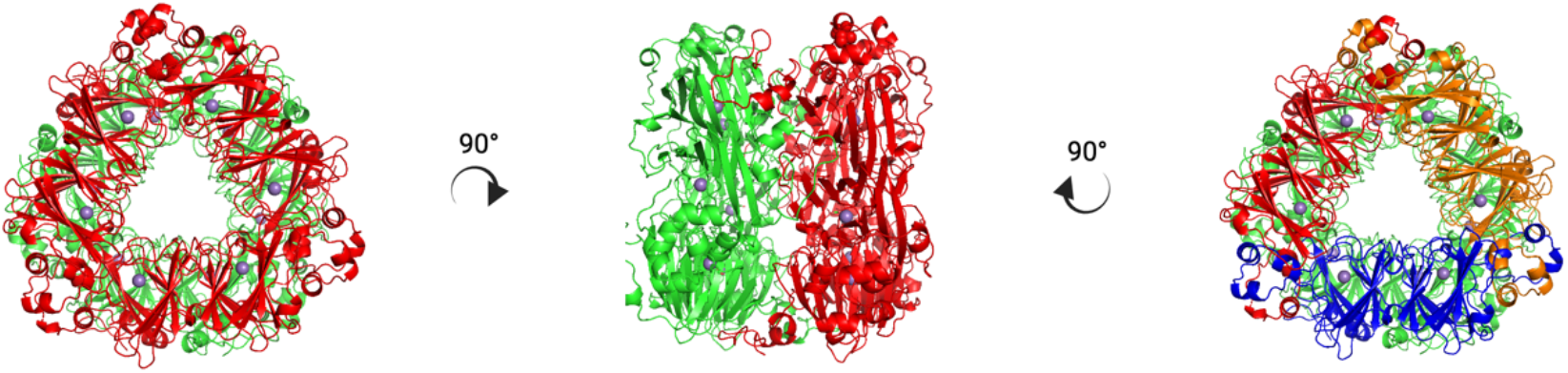
Structural features of *B. subtilis* Oxalate decarboxylase. Dimers of each trimeric unit are colored in red and green on the left and central image, while the three monomers forming the trimer are colored blue, orange and red, on the right image. The images have been created by using PyMOL Molecular Graphics System, Version 1.2r3pre, Schrödinger, LLC (22), starting from crystal crystal of OxDC using the PDB id: 1J58 (16).

Although the optimum catalytic activity of OxDC is observed at acidic pH (23), the biochemical characterization of OxDC under physiological conditions revealed the presence of a detectable activity at neutral pH, although associated with an increased tendency to unfold and aggregate. Thus, the therapeutic applications of OxDC are limited because they depend on activity under conditions of pH and ionic strength typical of a physiological environment.

In this work we have produced an engineered form of the enzyme more stable and active under physiological conditions as compared with the wild type counterpart. Specifically, guided by bioinformatic analyses and biochemical techniques, we have generated and purified a double mutant (I191F/H279D) showing enhanced *in vitro* thermal stability (Tm value 5°C higher), reduced propensity to aggregation under physiological conditions, and, more importantly, an increased catalytic activity at pH 7.2 (3-fold increase of the k_cat_/K_M_ value). By modelling approaches, we also defined the structural reasons behind the increased stability of the mutant. Overall, these data indicate that engineered OxDC is more efficient to degrade oxalate under physiological conditions, thus enhancing its potential as biological drug for genetic and non-genetic forms of hyperoxaluria.

## Results

### Rational engineering of OxDC using the “consensus-based approach”

Recent studies on the molecular features of *Bacillus subtilis* OxDC reported that at neutral pH the protein is relatively unstable and retains poor residual activity. A well-accepted and effective strategy to improve protein stability and activity is the “consensus-based” approach (24)^-^(25, 26). Consensus mutations are reversions of some protein residues to the ancestral amino acids, an approach that has been successfully applied to improve the overall stability and/or increase catalytic efficiency of several proteins (24, 26). Therefore, we generated the consensus sequence of *Bacillus subtilis* OxDC.

We selected a set of 11 mutations involving residues belonging to different protein domains (Figure 2A) based on the percentage of the consensus analysis (residues with the highest frequency at individual positions in the multiple sequence alignment used). From a structural point of view, mutation sites are spread over the entire dimer, being localized either on the monomer surface, in the monomer core, or in proximity of the monomer-monomer (dimer) interface of each trimer (Figure 2B).

**Figure 2.**
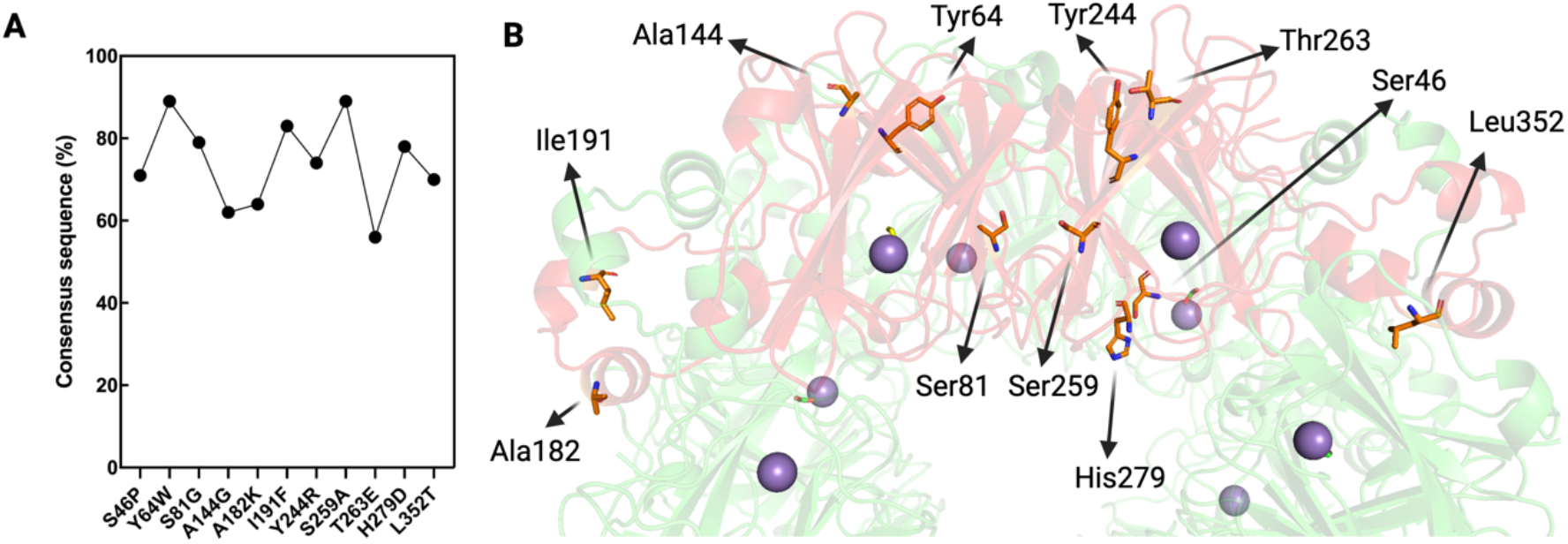
Consensus-based analysis of *Bacillus subtilis* OxDC. (**A)** Output of the consensus-based analysis obtained by using The ConSurf webserver. The graph shows the percentage of the selected amino acid substitutions on the OxDC sequence orthologues selected in this study. **(B)** Position of the amino acid residues selected as targets for mutagenesis within the OxDC structure (PDB id: 1J58). The monomer used to show the position of the consensus residues selected is highlighted and colored red, and each target residue is shown as orange sticks, while the remaining monomers are all colored green. The image has been created by using PyMOL Molecular Graphics System, Version 1.2r3pre, Schrödinger, LLC (27).

### Biochemical properties of single mutants

Each of the 11 selected mutations was introduced on the OxDC sequence by site-directed mutagenesis on the pET24a^(+)^-OxDC vector for bacterial expression (23). Seven protein variants were purified in high amounts from the soluble fraction and gave yields comparable to those of the wild type or even higher (Figure S1). On the other hand, the 4 variants Y64W, A144G, A182K, and L352T were present mainly in the insoluble fraction of the cellular lysate, suggesting that these mutations might affect the correct folding of the protein. For this reason, they were excluded from subsequent analyses.

The specific activity at different pHs and the melting temperature of the single mutants are shown in Figure 3. We found that the mutants I191F, Y244R, T263E and H279D display decarboxylase activity at pH 6.5 and 7.2 higher than OxDC wild type (Figure 3A). On the other hand, the mutations S46P, S81G and S259A do not increase the activity of OxDC at physiological pH. Analyzing the structural localization of the mutated residues, we noticed that beneficial mutations are mainly located on the protein surface (Y244R, T263E, and H279D) or the monomer-monomer interface (I191F). Notably, mutations involving residues located in the core of the OxDC monomer (such as Ser46, Ser81 and Ser259) do not increase activity at neutral pH and do not affect or even decrease thermal stability, thus indicating that alterations of the corresponding regions could possibly interfere with the proper folding of the monomeric subunits.

**Figure 3.**
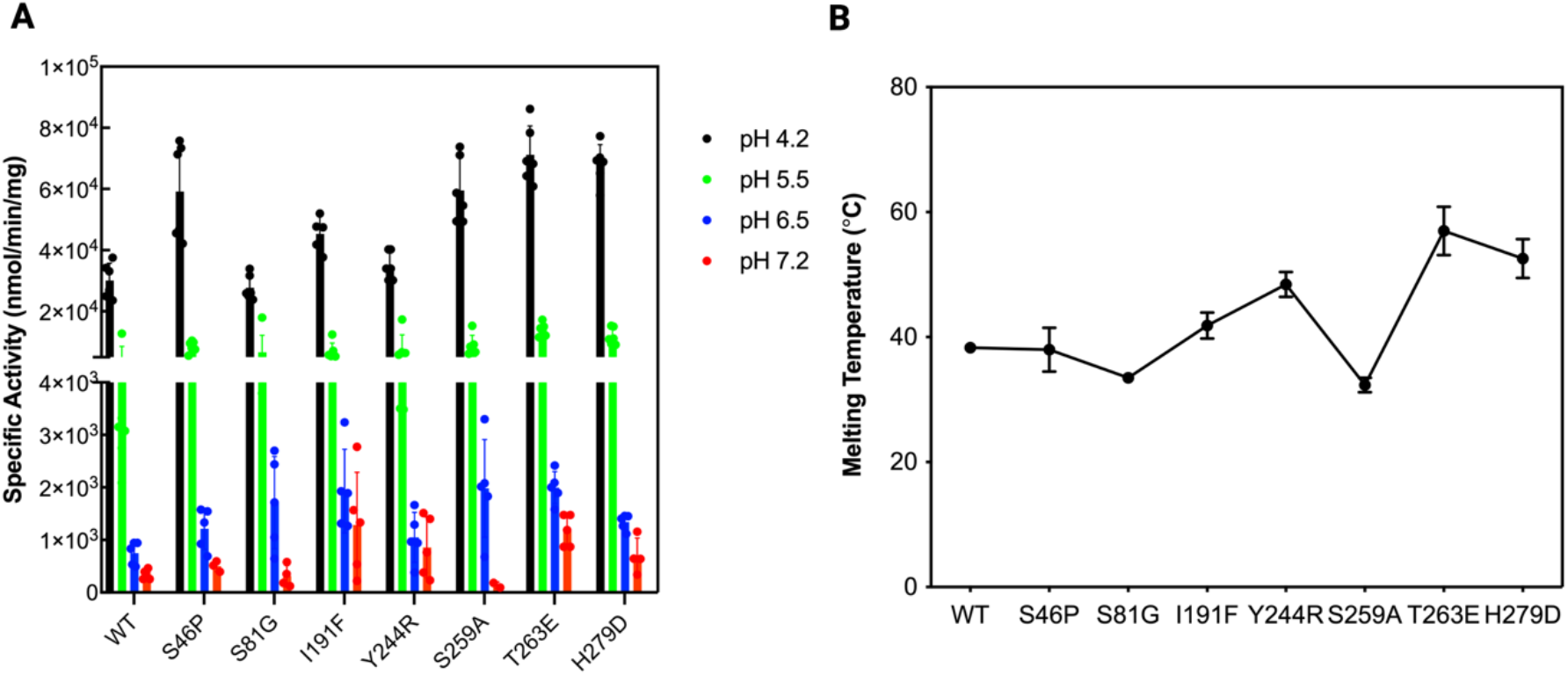
Functional and structural features of the selected single mutants. **(A)** Residual specific activity (expressed as nmol/min/mg) of OxDC wild type and of the single mutants, measured at different pHs (as indicated in the figure legend) at 25°C in 52mM sodium acetate pH 4.2, sodium acetate pH 5.5, 26 mM Bis-Tris pH 6.2 and PBS 1X pH 7.2 as previously reported here (23). **(B)** Plot of the values of thermal stability of OxDC wt and of the single mutants obtained by monitoring the changes of the CD signal at 222 nm in 16 mM Tris-HCl, 140 mM NaCl at pH 7.2.

### Generation and characterization of second-generation mutants by combinatory mutagenesis

We combined the four single beneficial mutations previously identified (I191F, Y244R, T263E and H279D), to produce the whole set of double and triple mutants as well as the quadruple mutant. We first evaluated the effects of combinatory mutagenesis on thermal stability and catalytic activity at saturating substrate concentration. Interestingly, as shown from the data in Figure 4A, the best results were obtained from double mutants, which show an increased melting temperature as compared to single, triple and the quadruple mutant. Specifically, the mutants I191F/T263E, I191F/H279D, Y244R/T263E and T263E/H279D show an increase in thermostability of 3-6 °C as compared to OxDC wild type. All of them also retain good catalytic activity at pH 7.2 except for the Y244R/T263E mutant.

**Figure 4.**
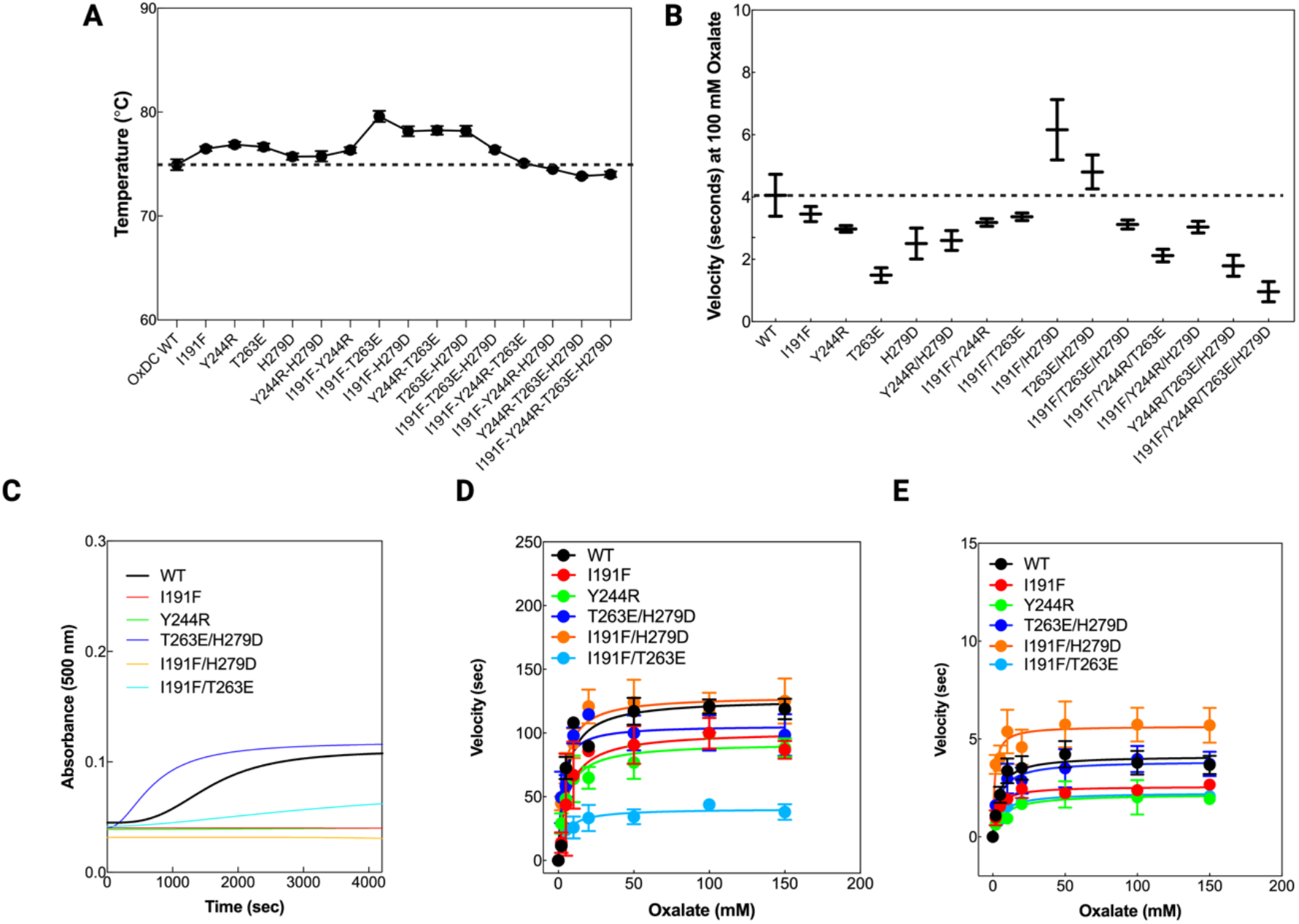
Structural and functional features of the selected OxDC mutants. **(A)** Experimental thermal shift values obtained by using the SYPRO orange probe. The dotted line indicates the OxDC wild type value of thermostability, used as reference. The experiments have been performed by using 1 μM protein and 5X SYPRO orange as final concentration in PBS 1X pH 7.2. Each experiment was repeated at least in triplicate for each mutant. **(B)** Turnover number expressed as velocity (seconds) using PBS 1X at pH 7.2 using 100 mM oxalate as substrate. **(C)** Turbidimetry experiments on the selected mutants performed by monitoring time-dependent changes in the absorbance at 500 nm. The experiments were performed by using 1 μM protein concentration at 37°C in PBS 1X. **(D)** Kinetic parameters of OxDC wild type and mutants measured by using 0.1μM protein concentration in sodium acetate 52 mM, NaCl 140 mM pH 4.2 at 37°C. The curves represent the fitting to the Michaelis-Menten equation. **(E)** Kinetic parameters of OxDC wild type and mutants measured by using 0.3 μM protein concentration in PBS 1X pH 7.2 at 37°C. The curves represent the fitting to the Michaelis-Menten equation. Data were collected and analysed in triplicate.

Based on the overall data obtained, we decided to further characterize the species showing an increase in melting temperature associated with an increased or preserved activity at pH 7.2: I191F, Y244R, T263E/H279D, I191F/H279D and I191F/T263E.

Since the two major concerns about OxDC wild type under physiological condition are related to its increased aggregation tendency and decreased catalytic efficiency, we focused on the stability and kinetic features of the selected mutants. Turbidimetry experiments revealed that OxDC wild type shows a considerable aggregation propensity at 37°C under conditions mimicking a physiological pH and ionic strength (PBS 1X pH 7.2) with a t_1/2_ = 1567 ± 21 seconds (Figure 4C). Interestingly, all the selected mutants showed slightly or no aggregation propensity under physiological conditions as compared to OxDC wild type, except the double mutant T263E/H279D that displays an increased aggregation rate (t_1/2_ = 699 ± 44 seconds).

By calculating the kinetic parameters of the mutants at pH 4.2, which represents the optimum of the enzyme, we confirmed that all retain detectable activity (Figure 4D), with values of catalytic efficiency for the decarboxylase reaction similar or lower to that of the wild type (Figure 4D and Table S2). However, the kinetic parameters of the mutated proteins under physiological conditions show a different scenario. At pH 7.2, most of the single and double mutants show a decreased catalytic efficiency compared to that of OxDC wild type, driven by both an altered k_*cat*_ and K_M_ value (Table S2).

On the other hand, the mutant I191F/H279D shows a catalytic efficiency increased by 3-fold. Based on the data obtained we pinpointed the I191F/H279D mutant as the best candidate, and we decided to characterize the structural features of the mutant I191F/H279D more in detail. We included in the structural analysis the two single mutants, I191F and H279D to dissect the contribution of each amino acid change to the structural properties of the double mutant. The comparison of intrinsic fluorescence spectra (which report on the microenvironment of the aromatic side chains) as well as of the ANS fluorescence spectra (which report on the presence of hydrophobic surfaces) provided evidence that the I191F/H279D mutant displays some conformational changes as compared with OxDC wild type. As shown in Figure 5A and 5B, the main alterations are related to (i) a consistent decrease (∼1.4-fold) of the maximum intensity in the intrinsic fluorescence spectra, and (ii) a ∼2-fold decrease in ANS emission fluorescence intensity along with a 4-nm red shift of the emission maximum.

**Figure 5.**
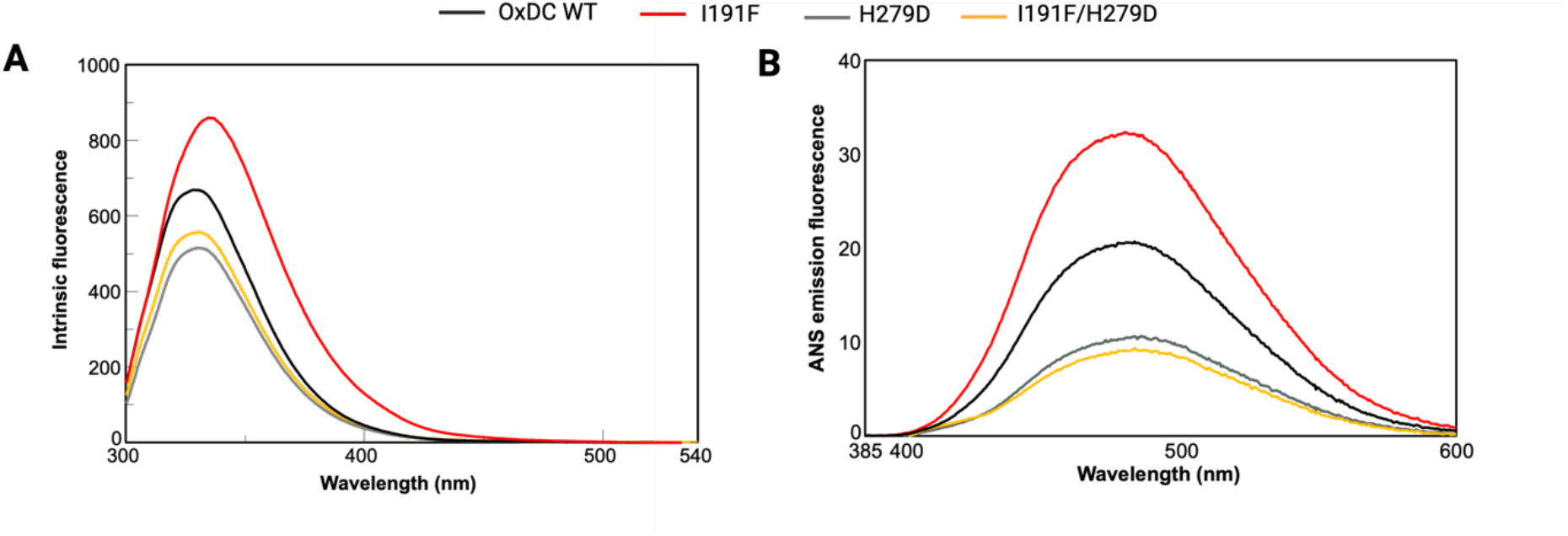
Spectral features of the double mutant I191F/H279D and the single mutants I191F and H279D compared to the OxDC wild type. **A**, Intrinsic emission fluorescence spectra of the selected mutants and OxDC wild type registered in PBS 1X using 0.25 μM protein concentration at 25°C. The λ_exc_ was 280 nm and the emission fluorescence recorded from 300 to 540 nm. **B**, ANS emission fluorescence spectra of the selected mutants and OxDC wild type registered in PBS 1X using 1 μM protein concentration at 25 C. The λ_exc_ was 365 nm and the emission fluorescence recorded from 380 to 600 nm.

In addition, by analyzing the spectroscopic data of the single mutants we noticed that the double mutant I191F/H279D shares similar spectroscopic features with the single mutant H279D whereas the mutation of Ile191 seems to induce larger conformational changes to the OxDC structure by itself. On the other hand, the kinetic parameters of the single mutants at pH 7.2 indicate that each of the I191F and H279D mutations (Table S2) alone do not increase the *k*_cat_ value or decrease the K_M_ value of the enzyme suggesting that some subtle changes occur at the active site of the double mutant probably as a consequence of a combined effect of the two mutations.

The finding that both the single mutations I191F and H279D mutations improve OxDC thermal stability (as previously discussed and reported in Figure 4A), although at a different extent, would suggest that the increased activity of the double mutant could be a secondary effect of the overall increased stability (in terms of increased thermodynamic stability and decreased aggregation propensity) under physiological conditions, possibly attributed to a different and more stable conformation of the mutated protein.

In order to test this hypothesis and have more deep insights into the stabilizing effects of the two mutations on the possible behavior of OxDC in a biological environment, we compared the OxDc wild type and double mutant for their sensitivity to proteolytic cleavage and to thermal stress.

The results shown in Figure 6 indicate that the double mutant I191F/H279D shows a slightly increased stability against the proteolytic cleavage mediated by the proteases chymotrypsin and pancreatin (Figure 6A and 6B), as well as an increased kinetic stability under thermal stress (Figure 6C). By fitting the residual activity of the proteins incubated at 37°C at different times, we found that the decay constant (k) of the double mutant is 31% lower as compared to the wild type counterpart. Taken together these results indicate that the double mutant shows an increased stability under temperature stress, due to a change in its structural features compared to the wild type counterpart.

**Figure 6.**
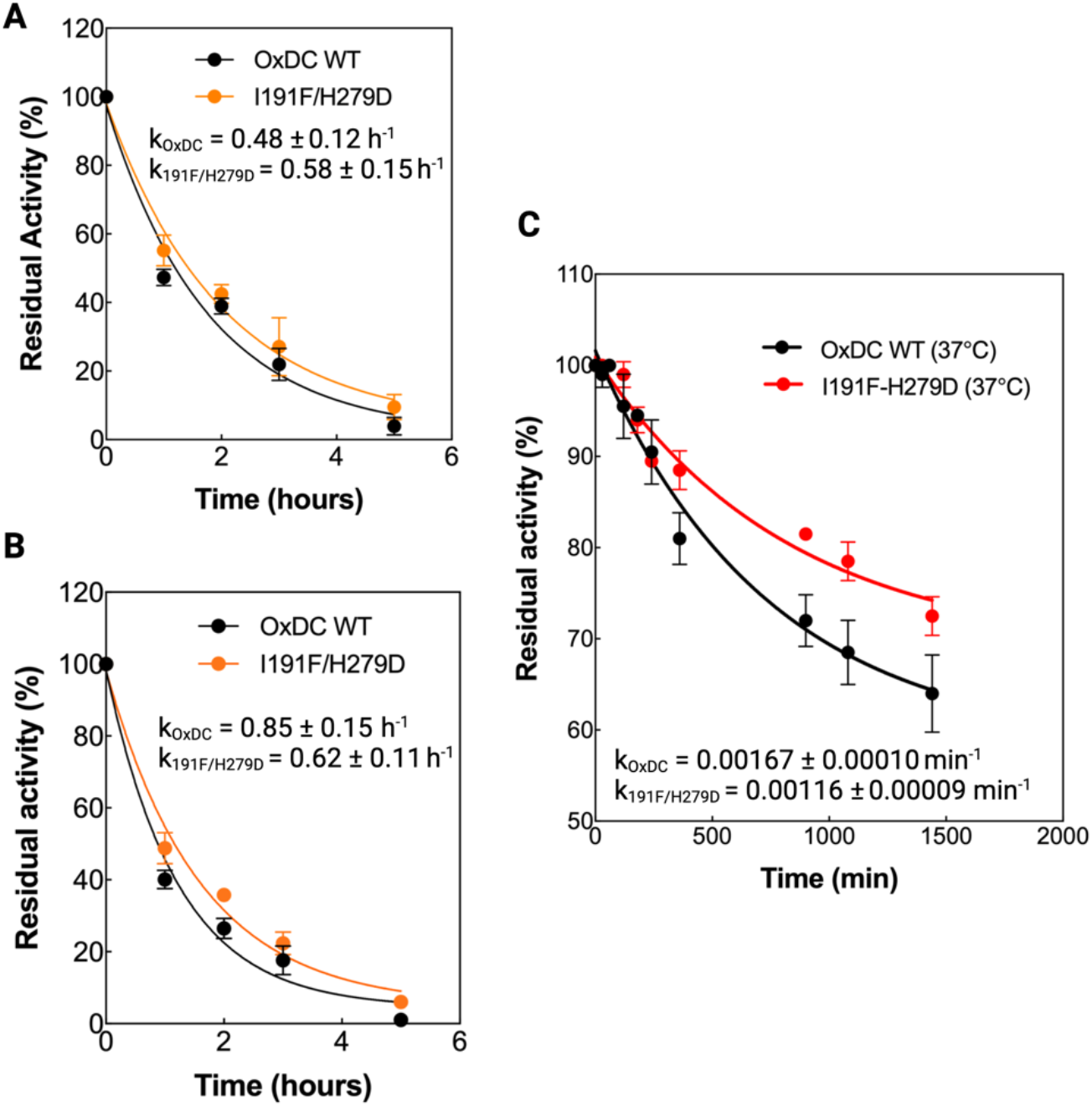
Residual activity plots in presence of two different proteases and under thermal stress at 37°C of OxDC WT and mutants I191F/H279D in PBS 1X pH 7.4. **(A) and (B)**, Residual activity expressed as % of OxDC WT and double mutants I191F/H279D in presence of pancreatin (A) and α-chymotrypsin (B) used at a ratio 50:1 for 5 hours at 37°C. **(C)** Kinetic stability under thermal stress of OxDC WT and the mutant I191F/H279D at 37°C at different incubation times (total time 24 hours). Within the graphs are reported the decay constants (k) obtained by fitting the experimental points using a single exponential decay function (see Material and Methods).

## DISCUSSION

OxDC is an enzyme endowed with a high potential for industrial applications, in particular for the treatment of hyperoxaluria, a group of pathologic conditions associated with increased oxalate excretion by either genetic or environmental causes, resulting in the formation of oxalate stones mainly in the kidneys (1, 3, 9). The consumption of high oxalate food represents an important risk factor for SH patients, who display increased intestinal oxalate absorption, but it is also discouraged in PH patients, who produce high amounts of endogenous oxalate (28). In addition, since dietary oxalate contributes to raise the amount of urinary oxalate thus promoting CaOx stones formation, a reduction in oxalate load can also affect the onset of more common kidney injuries, such as nephrolithiasis as well as acute and chronic kidney disease (29). As such, a wide number of potential patients could benefit from drugs able to reduce oxalate absorption at intestinal level. The oral administration of OxDC is one of therapeutic approaches proposed for the treatment of hyperoxaluria, to degrade intestinal oxalate thus preventing the intestinal absorption of food oxalate. Oral formulations of OxDC have been generated named Nephure™(30), OxDC CLEC(9), and Oxazyme. Nephure™(31) is used as a food ingredient, while OxDC CLEC and Oxazyme are cross-linked forms of OxDC from *B. subtilis* proven to be effective in healthy subjects even with limitations (9, 30, 31).

One of the main limitations related to the use of OxDC and others oxalate-degrading enzymes is the optimum pH of activity under acidic conditions (19, 32), even though the enzyme retains detectable decarboxylase activity at neutral pH (23). Nonetheless, under conditions mimicking the possible site of action as biological drug, OxDC shows a strongly reduced thermal stability associated with an enhanced propensity to unfolding and aggregation (23), which further compromise the efficiency in intestinal oxalate degradation.

Protein engineering studies are widely employed to improve catalytic efficiency and/or stability of enzymes employed as biological drugs or as tools for industrial applications (33, 34). One of the most popular strategies to increase the thermodynamic stability of a protein is the use of consensus-based approaches, which are based on multiple sequences alignments to introduce mutations that convert a low frequency amino acid with a more frequent one in a certain position (25, 26, 35).

Here we performed the engineering of OxDC from *B. subtilis* through a consensus-guided strategy. Upon a first screening of the effects of single selected mutations on the specific activity of the enzyme, we chose the favorable ones and started a combinatorial mutagenesis study to foster protein stability by exploiting possible non-additive effects of multiple mutations (36, 37). We obtained the I191F/H279D double mutant, which shows an increased catalytic activity at neutral pH, increased thermodynamic stability and almost undetectable propensity to aggregation under physiological condition of ionic strength and pH. These effects are similar to those obtained on other enzymes using analogous strategies (24, 38–42). A comparative analysis of the effects of each single mutation indicates that both increase the melting temperature. The crystal structure of OxDC indicates that both Ile191 and His279 are far from the active site. Ile191 is located in an alpha-helix of the cupin domain I in a region that forms a claw-like protrusion which is directly involved in subunits contacts (Figure 7A). Ile191 is involved in a hydrophobic cluster formed by Ile180, Ala181, Val186, Ile194 and with Leu361, Leu363, Phe367 and Leu371 of the neighboring subunit. This hydrophobic cluster stabilizes the region which is of fundamental importance for OxDC quaternary structure. In fact, the trimeric layers of the hexamer are stabilized by the interlocking claw-like α-helical protrusions of adjacent monomers (16, 20).

**Figure 7.**
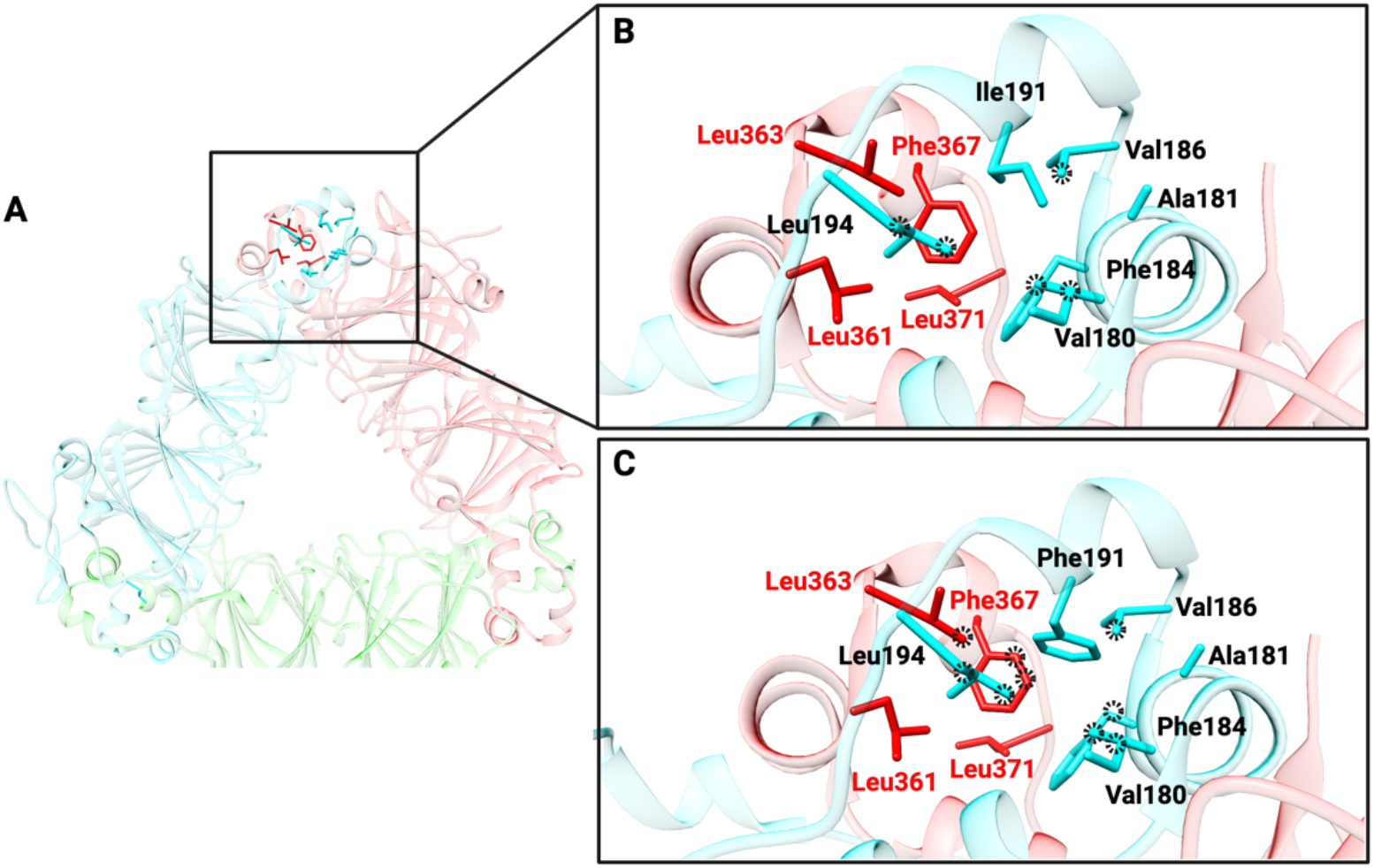
In silico analysis of the effects of the I191F mutation. **(A)** Localization of the residue Ile191 in the OxDC crystal structure. The OxDC dimers are colored in red, cyan and green. **(B)** Zoom-in of the interactions mediated by Ile191 (cyan) with the surrounding residues. The interactions are highlighted using black dot circles. **(C)** Molecular modeling of the mutation Ile191Phe and analysis of the interactions mediated by Phe191 with the surrounding residues. The mutant I191F has been prepared by using the tool “rotamers” (43) present in the program UCSF Chimera X-1.7 (44).

Interestingly, Ile191 side chain mainly interacts with residues located on the same subunits, as reported in Figure 7B. However, the substitution of Ile in position 191 with a bulky Phe residue seems to increase the total number of interactions, especially those between the two subunits of the trimer (Figure 7C) without generating steric hindrance with the surrounding environment. In detail, Phe191 directly interacts with Leu363 and Phe367, two residues located on the adjacent subunits, playing an important role in stabilizing the trimeric layers of the OxDC hexamer, and possibly explaining the increased thermostability of the single mutant.

On the other hand, residue His279 is located in the cupin domain II and it is exposed to the solvent, coordinating solvent molecule and mediating a hydrogen bond with Phe315 mainchain (16). As reported in Figure 8A, the electrostatic potential map of OxDC wt surface at pH 7.0 shows interesting chemical-physical features: a large negatively charged patch on one side of the surface along with a large uncharged patch on the other side. The same analysis on the H279D mutant (Figure 8B) reveals that the presence of the three Asp decreases the area of the not-charged patches along the surface, possibly explaining the low signal of the probe ANS reported in Figure 5B, which indicates a decreased presence of hydrophobic surfaces.

**Figure 8.**
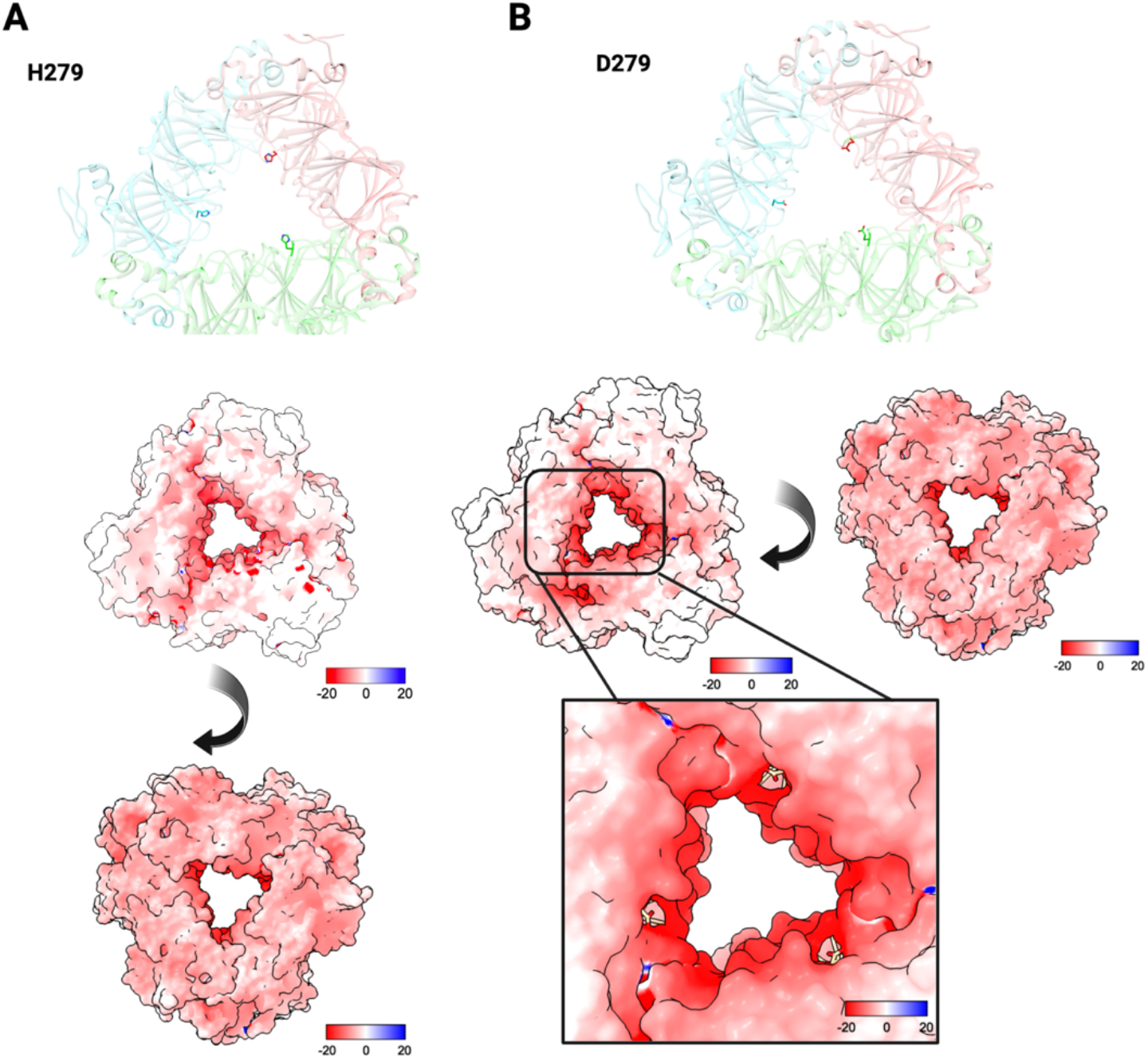
Electrostatic potential maps of OxDC WT and the H279D mutant. **(A)** Location of the His279 residues in the crystal structure and electrostatic potential map of OxDC WT. **(B)** Electrostatic potential map of mutant H279D. The mutant has been prepared using the tool “Rotamers” present in UCSF Chimera X-1.7 (44). The electrostatic maps have been calculated by using the software suite APBS-PDB2PQR at pH 7.2 (https://server.poissonboltzmann.org/pdb2pqr), at 37°C and visualized using UCSF Chimera X-1.7 (44).

In this regard, it is of note that also the double mutant shows a reduced exposure of hydrophobic surfaces with a concomitant increase of negatively charged surfaces, and that the H279D mutation is probably the main contributor to this effect. This can provide a possible explanation to the reduced aggregation tendency under physiological conditions shown in Figure 4C.

More difficult is the interpretation of kinetic data, because the I191F/H279D double mutant displays an increased catalytic efficiency despite none of the single mutations specifically affects the *k*_cat_ and/or the K_M_ of OxDC. It can be suggested that long-range interactions are responsible for the increased catalytic efficiency, as already reported for other enzymes (45)^-^ (46, 47). In addition, the increased activity of the double mutant at pH 7.2 can be a secondary effect of the increased half-life of the protein under physiological conditions (Figure 6C).

Overall, although in silico analyses provide a structural explanation to the effects observed in vitro, it must be taken into account that the overall thermodynamic stability of enzymes is dependent on several intrinsic features of the proteins. Indeed, the flexibility of its component stretches of amino acid, long-range interactions such as networks of hydrogen bonds and increased number of hydrophobic interaction, can influence the overall resistance to external stresses, finally resulting into a different half-life in cellular and non-cellular systems (46–48).

In conclusion, we obtained an engineered form of OxDC from *B. subtilis*, showing improved properties in terms of activity and stability at physiological conditions. Our data could help in ameliorating currently available drugs based on OxDC administration, although studies in animal models will be necessary to assess the effects of engineering on the *in vivo* performance of the enzyme.

### Experimental Procedures

#### Materials

Potassium oxalate, sodium formate, potassium permanganate, isopropyl-β-D-thiogalactopyranoside, manganese chloride and imidazole were purchased from Merck Life Science srl (St. Louis, MO, USA). Oligonucleotides for site directed mutagenesis were purchased from Bio-Fab Research srl (Rome, Italy). All other chemicals used were of analytical grade.

#### Consensus-based approach

The Consensus based sequence has been calculated using the ConSurf web server (http://consurf.tau.ac.il/2016/) (49). The homologues sequences of OxDC were collected from UNIREF90 database and selected by using HMMER (E-value cut-off was 0.0001 and number of iterations 1). The multiple sequence alignment of 150 homologues sequences (identity from 95 % to 35%) has been performed by using the MUSCLE (Multiple Sequence Comparison by Log-Expectation) (50). The consensus sequence-based alignment has been visualized and analyzed using Jalview 2.10.5 (https://www.jalview.org) (51).

#### Molecular Modeling and electrostatic potential maps

Visual inspection of the oxalate decarboxylase crystal structure (PDB id: 1J58) was performed using the software Python-enhanced Molecular Graphics tool (PyMol) (Schrödinger, LLC) (22) and UCSF Chimera X-1.7 (National Institute of Health, NIH) (44) The dimeric structure was obtained using the Proteins, Interfaces, Structures and Assemblies (PISA) PDBePISA web server (52), starting from the available coordinate file of the monomer (PDB id:1J58). The mutant I191F and H279D have been prepared using UCSF Chimera X-1.7 (44) with the rotamers tool using the Dunbrack rotamer library (43). The electrostatic maps of the OxDC WT and mutant H279D have been calculated by using the webserver APBS-PDB2PQR software suite (https://server.poissonboltzmann.org) (53). In detail, the analysis of the protonation states of the residues at pH 7.0 were carried out using the PDB2PQR suite tool (using CHARMM as forcefield) which create an input file for the calculation of the electrostatic potential map using APBS suite. The graphic visualization of the electrostatic potential maps of the OxDC wild type and H279D mutant have been performed using UCSF Chimera X-1.7 (44).

#### Site-Directed mutagenesis

The selected mutations have been introduced by site-directed mutagenesis using the QuikChange II site-directed mutagenesis kit (Stratagene San Diego, California) using the vector encoding the sequence of *Bacillus subtilis* OxDC (Uniprot ID: O34714) cloned in the vector (pET24a-OxDC). The oligonucleotides used for the mutagenesis are listed in Table S1. All the mutations were confirmed by the entire DNA sequencing.

#### Expression and Purification of OxDc and mutants

His-tagged OxDC wild type and the selected mutants were expressed in *E. coli* and purified by affinity chromatography with minor modifications from the previous protocol (23).

In detail, *E. coli* BL21 (DE3) cells transformed with the constructs pET24a-OxDC were grown in Luria Broth (LB) in a total volume of 0.5 litres at 37 °C. Cells were grown with vigorous shaking to an OD of 0.3-0.4 at 600 nm 5 mM MnCl_2_ was added to the culture and protein expression was induced with 0.2 mM IPTG for 16 hours at 30°C. Cells were then harvested by centrifugation and resuspended in lysis buffer (50 mM Tris-HCl pH 8 containing 0.5 M NaCl, 20 mM imidazole and EDTA-free protease inhibitor cocktail). After sonication, cell debris were removed by centrifugation (16000 x *g* for 30 minutes at 4°C). The lysate was loaded on a homemade Nichel-Affinity resin column (2 mL of resin) equilibrated with 50 mM Tris-HCl pH 8 containing 500 mM NaCl and 50 mM imidazole. At this point, 10 mL of the same buffer containing 500 mM imidazole was applied. Upon forced dialysis by Amicon ultra 10 devices (10 kDa) and wash with storage buffer (50 mM Tris-HCl pH 8, 500 mM NaCl and 5 mM DTT), OxDC-His wild type and the mutants have been conserved at -20°C. The purification was evaluated by SDS-PAGE: 1 μg of OxDC was loaded per lane on 10% SDS-PAGE (Figure S1A, B). The protein concentration was measured by using ε_280_ = 42,340 M^-1^ cm^-1^ and the yield is reported in Figure S1C.

#### Spectroscopic measurements

Absorption measurements for protein quantification were carried out using a Jasco V-750 spectrophotometer with 1-cm path length quartz cuvettes (Jasco Europe S.r.l.). 8-anilino-1-naphthalene sulfonate (ANS) fluorescence emission spectra were recorded on a Jasco FP8200 spectrofluorometer equipped with a thermostatically controlled cell holder by using 1 cm path length quartz cuvettes. Excitation was set at 365 nm with both the excitation and the emission slits of 5 nm. The ANS-binding experiments have been performed at 25°C using 1 μM protein concentration in 16 mM Tris-HCL, 140 mM NaCl, pH 8.0. The increase in turbidity was monitored by measuring the absorbance at 500 nm as a function of time using a MultiSkan SkyHigh microplate reader (Thermo Fisher Scientific) at 37°C using 1X PBS buffer in presence of 1 μM protein concentration in a final volume of 200 μL.

#### Enzymatic activity measurements

OxDc wild type and mutants’ activity at different pHs has been measured by detecting the product formate using potassium permanganate. Potassium permanganate is a chemiluminescent reagent often used to detect organic molecules(13). It has absorption maxima at three different wavelengths, specifically at 525, 545 and 569 nm. It has been reported(54) that the addition of 1 mM potassium permanganate solution leads to a significant decrease of absorbance maxima in presence of formate. Here, after an initial setup of the measurements in presence of different formate concentrations and at fixed potassium permanganate concentration of 1 mM, we decided to use the 545 nm to detect the formate produced by the enzymatic reaction. In detail, the OxDc enzymatic reaction has been performed at two different pHs. At pH 4.2 we have used the sodium acetate 52 mM in presence of NaCl 140 mM, using 0.1 μM OxDc (wild type and mutants) at 37°C for 10 minutes in presence of different concentrations (0-150 mM) of potassium oxalate. At pH 7.2 we have used PBS 1X using 0.3-0.5 μM OxDc (wild type and mutants) at 37°C for 25 minutes in presence of different concentrations (0-150 mM) of potassium oxalate. The reaction mix have been stopped by using KP 100 mM pH 8.0 in presence of 1 mM potassium permanganate (final concentration) in a total volume of 200 μL and the absorbance at 545 nm of the first 300 seconds has been measured by using MultiSkan SkyHigh microplate reader at 25°C and analysed using GraphPad Prism 10.2.1 (GraphPad Software, Boston, Massachusetts USA, www.graphpad.com). The calibration curves used to analyzed the data is reported in Figure S2.

#### Thermal shift assay

Thermal shift assay was used to estimate and compare the thermal stability of OxDC wild type and mutants using a high-throughput differential scanning fluorimetry (DSF) assay. We used SYPRO orange 5000x (Sigma Aldrich) as fluorescent dye. The fluorescence of the dye is quenched in aqueous solution while it can bind exposed hydrophobic regions upon protein unfolding. We used a final concentration of 5x SYPRO orange dye (1:1000 v/v) with 1 μM protein in 20 μL as total volume. The experiments were performed using the StepOnePlus real-time instrument (Thermo Fisher Scientific) in a 96-well plate. The data were analyzed using a script and repeated for at least 3 times using the same experimental conditions.

#### Proteolysis experiment

0.5 mg of OxDC WT and I191F/H279D have been resuspended in PBS 1x pH 7.2 along with 10 μg of proteases pancreatin and α-chymotrypsin (ratio 50:1 between OxDC and the proteases). The reaction mix have been incubated for 5 hours at 37°C and at each time point 100 μL were withdrawn, and the activity tested by using the assay reported in the paragraph “Enzymatic activity measurements” by using potassium permanganate. The substrate concentration used to test OxDC enzymatic activity was 100 mM oxalate. The *t*_1/2_ and the kd values were calculated using a single exponential decay equation using GraphPad Prism 10.2.1.

#### -Determination of the kinetic stability of the OxDC WT and mutant I191F/H279D

To determine the kinetic stability of the OxDC WT and the mutant I191F/H279D we have incubated the enzymes at 37°C in PBS 1X pH 7.2, using 1 μM protein concentration, and measuring the residual activity at different incubation time. For all the measurements, the initial activity of the proteins (with no incubation) was defined as 100% and the subsequent points at the different incubation times were calculated (%) using as reference the unincubated point. The *t*_1/2_ and the k values were calculated using a single exponential decay equation using GraphPad Prism 10.2.1 (GraphPad Software, Boston, Massachusetts USA, www.graphpad.com).

#### Statistical analysis

Data have been analyzed using GraphPad Prism 10.2.1 (GraphPad Software, Boston, Massachusetts USA, www.graphpad.com). All the data presented in this work represent the mean ± SD of at least two or three independent experiments.

## Data Availability

All data and materials are available upon reasonable request.

## Funding and Additional Information

The study was supported by the Italian Ministry of University and Research (SIR project RBSI148BK3) to B.C.. P.L was supported by the Okinawa Institute of Science and Technology Graduate University (OIST) with subsidy funding from the Cabinet Office, Government of Japan. M.D. thanks the financial support from Japan Society for the Promotion of Science (JSPS) for the Kakenhi Early Career Scientist N. 22K15065. P.L. thanks the financial support from Takeda Grant.

## Acknowledgements

We thank Dr. Ben E. Clifton for the suggestions and discussion on the project and the help for the set-up and data analysis of the thermal shift assay. The images of the present work were prepared using Biorender.com.

