## Supplementary Information for "Engineered Oxalate decarboxylase boosts activity and stability for biological applications"

[1]Department of Medicine and Surgery, Section of Physiology and Biochemistry, University of Perugia, 06132, Perugia, Italy
[2]Protein Engineering and Evolution Unit, Okinawa Institute of Science and Technology (OIST), Onna, Okinawa 904-0495, Japan
[3]Center of Cooperative Research in Biosciences (CIC bioGUNE) Basque Research and Technology Alliance (BRTA), Bizkaia, Technology Park, 48160 Derio, Spain
[4]Department de Quimica Fisica, Unidad de Excelencia en Quimica Aplicada a Biomedicina y Medioambiente e Instituto de Biotecnologia, Universidad de Granada, Granada, Spain
[5]Department of Biomolecular Sciences, University of Urbino "Carlo Bo", Urbino, Italy
[6]Institute of protein Research, Osaka University, Suita, Osaka 565-0871, Japan

**Table S1. Primer sequences used for mutagenesis of OxDC.**

| | Forward primer sequence | Reverse primer sequence |
|---|---|---|
| S46P | 5'-taaagctgaatttcatgttgggaacggtaccatggtcggtc-3' | 5'-gaccgaccatggtaccgttcccaacatgaaattcagcttta-3' |
| S81G | 5'-acgcatgttaacgcccgccaggttctcgc-3' | 5'-gcgagaacctggcgggcgttaacatgcgt-3' |
| I191F | 5'-ccggcaggttgctgaactcctctttggtaac-3' | 5'-gttaccaaagaggagttcagcaacctgccgg-3’ |
| Y244R | 5'-tggtgctatccgcaatacgcaccttgccaccttcgc-3' | 5'-gcgaaggtggcaaggtgcgtattgcggatagcacca-3' |
| S259A | 5'-cggtcaccagcgcggccgcaatggtcttgc-3' | 5'-gcaagaccattgcggccgcgctggtgaccg-3' |
| T263E | 5'-cgcacccggttcaacctccaccagcgcgctcgc-3' | 5'-gcgagcgcgctggtggaggttgaaccgggtgcg-3' |
| H279D | 5'-gtactgccattcgtcggtgttcgggtgcc-3' | 5'-ggcacccgaacaccgacgaatggcagtac-3' |

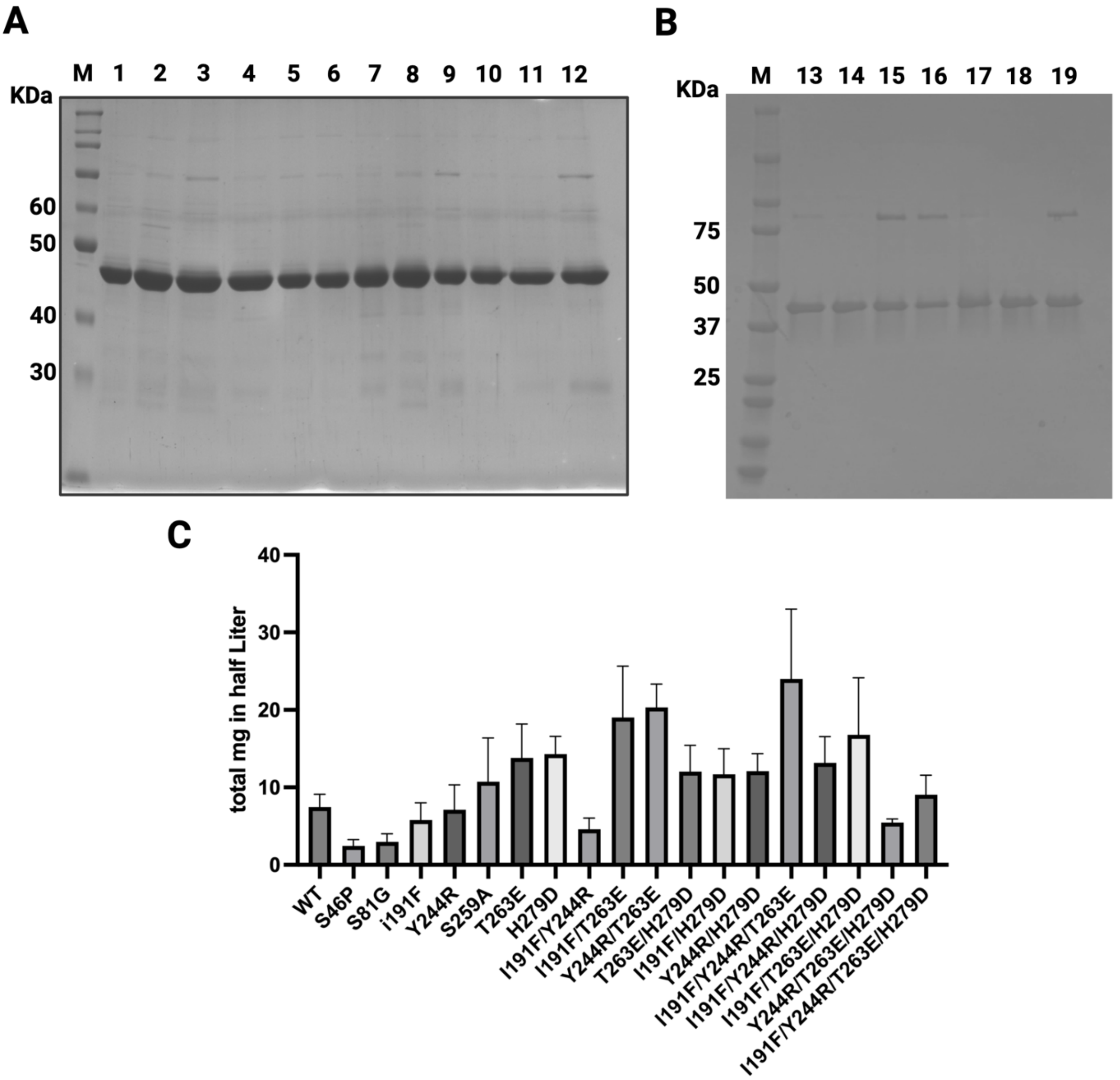


**Figure S1. SDS–PAGE analysis of purified recombinant OxDC variants and yield of the purification**. **Panel A and B:** SDS-PAGE gel of the purified proteins involved in the study. Lane M, protein marker, from 1 to 19 the purified proteins. In detail: 1: WT; 2: S46P; 3: S81G; 4: I191F; 5: Y244R; 6: S259S; 7: T263E; 8: H279D; 9: I191F/Y244R; 10: I191F/T263E; 11: Y244R/T263E; 12: T263E/H279D; 13: I191F/H279D; 14: Y244R/H279D; 15: I191F/Y244R/T263E; 16: I191F/Y244R/H279D; 17: I191F/T263E/H279D; 18: Y244R/T263E/H279D; 19: I191F/Y244R/T263E/H279D. **Panel C:** yield expressed as total milligrams obtained from the purification of the proteins involved in this study.

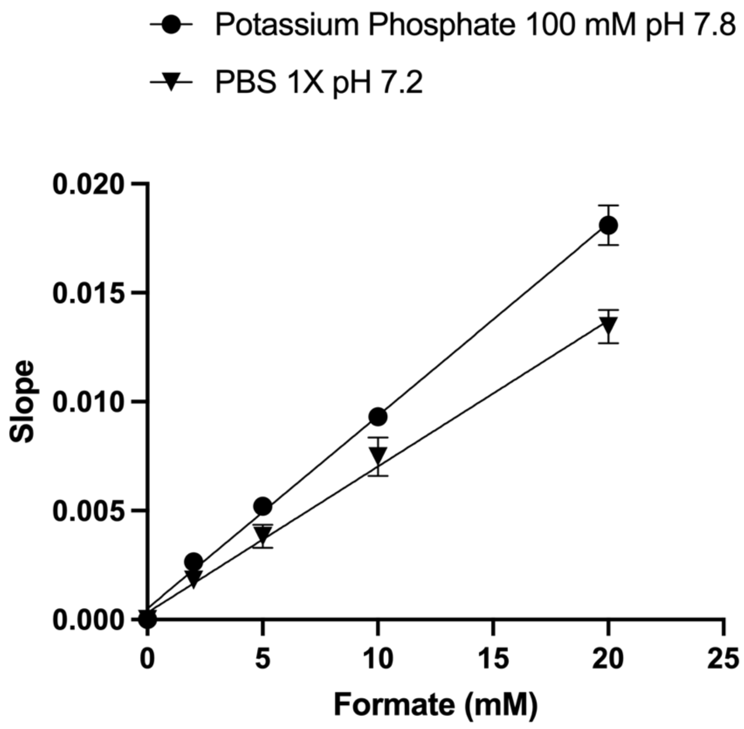


**Figure S2**. **Assay and calibration curve used for the detection of formate using potassium permanganate in KP 0.1 M pH 7.8 and PBS 1X pH 7.2.**

**Table S2**. **Kinetic parameters of OxDC wild type and mutated proteins obtained using sodium acetate 52 mM, NaCl 140 mM pH 4.2 or PBS 1X pH 7.2 at 37°C.**

| | **Protein** | **$k_{cat}$ ($s^{-1}$)** | **$K_M$ (mM)** | **$k_{cat}/K_M$ ($s^{-1}mM^{-1}$)** |
|---|---|---|---|---|
| **pH 4.2** | | | | |
| | Wild type | 126 ± 17 | 5 ± 2 | 25.2 ± 13.5 |
| | I191F | 101 ± 15 | 6 ± 2 | 16.8 ± 8.0 |
| | Y244R | 93 ± 10 | 5 ± 1 | 18.6 ± 5.7 |
| | H279D | 49 ± 7 | 6 ± 2 | 8.2 ± 2.9 |
| | I191F/T263E | 41 ± 5 | 4 ± 1 | 10.2 ± 3.8 |
| | I191F/H279D | 129 ± 11 | 4 ± 2 | 32.2 ± 18.8 |
| | T263E/H279D | 106 ± 12 | 3 ± 1 | 35.3 ± 15.7 |
| **pH 7.2** | | | | |
| | Wild type | 3.6 ± 0.6 | 3.1 ± 1.2 | 1.16 ± 0.47 |
| | I191F | 3.6 ± 0.3 | 2.4 ± 0.9 | 1.50 ± 0.43 |
| | Y244R | 2.2 ± 0.5 | 7.0 ± 2.4 | 0.31 ± 0.11 |
| | H279D | 2.2 ± 0.4 | 6.0 ± 2.2 | 0.36 ± 0.14 |
| | I191F/T263E | 2.0 ± 0.2 | 5.2 ± 2.0 | 0.38 ± 0.13 |
| | I191F/H279D | 5.5 ± 0.9 | 1.8 ± 0.5 | 3.05 ± 1.23 |
| | T263E/H279D | 4.9 ± 0.5 | 3.6 ± 1.1 | 0.57 ± 0.17 |
